# Biased re-orientation in the chemotaxis of peritrichous bacteria

**DOI:** 10.1101/2020.11.11.379230

**Authors:** T. Nakai, T. Ando, T Goto

## Abstract

Many kinds of peritrichous bacteria that repeat runs and tumbles by using multiple flagella exhibit chemotaxis by sensing a difference in the concentration of the attractant or repellent between two adjacent time points. If a cell senses that the concentration of an attractant has increased, their flagellar motors decrease the switching frequency from counterclockwise to clockwise direction of rotation, which causes a longer run in swimming up the concentration gradient than swimming down. We investigated the turn angle in tumbles of peritrichous bacteria swimming across the concentration gradient of a chemoattractant because the change in the switching frequency in the rotational direction may affect the way tumbles. We tracked several hundreds of runs and tumbles of single *Salmonella typhimurium* cells in the concentration gradient of L-serine, and found that the turn angle depends on the concentration gradient that the cell senses just before the tumble. The turn angle is biased toward a smaller value when the cells swim up the concentration gradient, whereas the distribution of the angle is almost uniform (random direction) when the cells swim down the gradient. The effect of the observed bias in the turn angle on the degree of chemotaxis was investigated by random walk simulation. In the concentration field where attractants diffuse concentrically from the point source, we found that this angular distribution clearly affects the reduction of the mean square displacement of the cell that has started at the attractant source, that is, the bias in the turn angle distribution contributes to chemotaxis in peritrichous bacteria.

**Statement of Significance:** We found another aspect in the chemotactic behavior of peritrichous bacteria. Chemotactic behaviors in peritrichous bacteria were predicted to be observed at the turn angles during tumbles motion as well as at the duration of runs; smaller changes in the swimming direction of cells swimming up the attractant’s gradient can be observed. This behavior is appropriate because the nature of bacterial chemotaxis changes the switching rate of the rotational direction of the flagellar motors according to the environment. Cells swimming upward reduce the turn angle by switching fewer flagellar motors to loosen flagellar filaments from the bundle during tumbles. We have shown that this prediction is correct.

## I. Introduction

Many kinds of bacteria exhibit chemotaxis; the cells accumulate around a favorable substance (attractant) or recede from an unfavorable one (repellent). The mechanism of chemotaxis has been intensely studied in peritrichous bacteria [1–3], such as *Escherichia coli* and *Salmonella typhimurium*, which have multiple flagella. Cells swim by rotating the helical flagella with the help of flagellar motors located at the proximal ends of the flagella. When each motor rotates counterclockwise (CCW), when observed from the distal end of the flagellum to the motor, the flagella form a bundle and the cell propels itself (run). When the rotational direction of the motors switches to clockwise (CW), the corresponding flagella are released from the bundle and the swimming direction of the cell changes (tumble) [4–6]. For *E. coli* cells, the mean time durations of run and tumble movements are approximately 1 and 0.1 s, respectively [7].

Cells exhibit chemotaxis upon sensing a difference in the concentration of the attractant or repellent between two adjacent time points [8, 9]. If a cell senses that the concentration of an attractant has increased, their flagellar motors decrease the switching frequency from CCW to CW direction of rotation, which causes a longer run in swimming up the concentration gradient than swimming down. Though a cell cannot move in a particular direction after exhibiting tumble motion or situate itself at the optimal position (with the maximum concentration of the attractant), the cell can remain at region where the concentration of the attractant is comparatively high. Longer runs toward the attractant have been observed in *E. coli* [7,9] and *Salmonella typhimurium* [10].

Recently, Sourjik et al. predicted that chemotaxis, in peritrichous bacteria, can be observed at the turn angles during tumbles motion [8]. The authors in the article refer to smaller changes in the swimming direction of cells swimming up the attractant’s gradient. This prediction is based on an observational study with regard to flagellar filaments in *E. coli* cells during tumble, which indicated that the turn angle reduces when fewer filaments are released during a tumble [11]. *E. coli cells* have 5-6 flagella on an average, and the corresponding flagellar motors can independently switch the direction of rotation [12]. The number of flagella released during tumble varies and can decrease when the switching from CCW to CW direction of rotation is suppressed when a chemoattractant is sensed. Turner et al. also demonstrated that the turn angle distribution is uniform (random direction) when many flagella are released during a tumble. The apparent influence of this change in the turn angle distribution on the chemotactic behavior has been confirmed by performing numerical simulation [13]. Experimentally, Saragosti et al. demonstrated that the drift velocity of *E. coli* cells in the attractant gradient has to be estimated by biases in the distribution of the turn angles as well as the duration of runs [14].

Our aim, in this study, was to demonstrate that the turn angle distribution depends on the swimming direction (up or down the concentration gradient). We observed the chemotactic behavior of a single cell for a long time to directly relate the cell’s movement with the degree of accumulation. Short-time observation of many cells would result in obtaining ambiguous information about characteristic motions because of difference in the swimming motion of each cell; hence, we chose to observe for a longer period of time.

## II. Methods

A strain of peritrichous bacteria, *Salmonella typhimurium*, SJW1103, was used. Cells were cultured in 3 mL of Luria-Bertani (LB) medium (1% polypeptone, 0.5% yeast extract, and 0.5% sodium chloride) overnight at 30°C. Then, 0.1 mL of the bacterial suspension was added to 3 mL of the motility buffer (pH 7.0, 10 mM KH_2_PO_4_, 0.1 mM EDTA, and 10 mM sodium lactate) and incubated for 3 h at 30°C. The solution was centrifuged for 1 min at 10000 rpm (CHIBITAN-II, Merck, Japan). The supernatant 0.9 mL was removed, and same amount of buffer was added to remove the culture medium for the detection of an attractant. The size of the cells was 1–2 μm in length and 0.6–0.8 μm in width.

Capillary assay was performed, in which cells were made to accumulate around the tip of a capillary filled with an attractant (same as that followed by authors in Ref. 10). A capillary was made by pulling a glass tube with a filament (GD-1, Narishige, Japan) using a puller (PC-10, Narishige, Japan). The inner diameter of the tip was 3–4 μm. L-serine (MW = 105; Wako, Japan) was used as the attractant. To prevent the flow as a result of capillary force, the attractant solution was jellified with agar. One molar L-serine with 0.3% agar (Agarose L, Wako, Japan), dissolved in the motility buffer, was heated with the help of a hot plate at 80°C and then aspirated into the capillary. Then, the outer surface of the capillary was washed with distilled water. As shown in Fig. 1, the capillary and the bacterial suspension were inserted between the slide glass (S7224, 76 mm × 26 mm, thickness 1.2–1.5 mm, Matsunami, Japan) and cover glass (18 mm × 18 mm, thickness 0.13–0.17 mm, Matsunami, Japan). Bacteria were aspirated using a microinjection (IM-6, Narishige, Japan) with a capillary (inner diameter of approximately 100 μm) and poured into the preparation. Then, 150 μL of buffer was added. To prevent convection as a result of evaporation, the edge of the cover glass was sealed with petroleum jelly. The thickness of the suspension was approximately 300 μm.

**Fig. 1.**
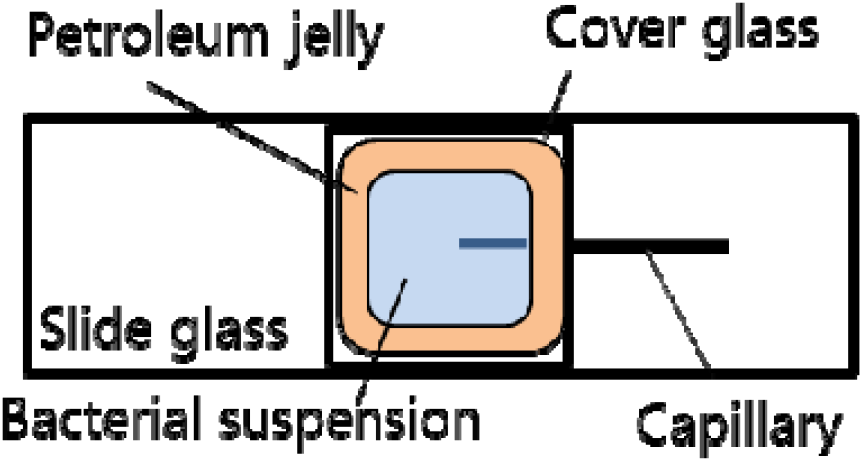
Top view of the prepared slide for observation of the bacterial chemotaxis.

Cells accumulating around an attractant were observed using an inverted microscope (IX71, Olympus, Japan) with an objective lens (LUCPLFLN40XPH, NA = 0.60, Olympus, Japan). The capillary tip was set at the center of the view. Images were recorded using a digital camera (DP27, Olympus, Japan) at 30 fps with a resolution of 1216 × 960 pixels (corresponding to the field of view of 280 × 220 μm). Images were analyzed using a tracking software (DippMotionPro, Ditect, Japan) and the two-dimensional trajectories of cells were measured.

As the flagella were not visible in bright field observation, the time of tumble was judged from the change in both the swimming direction and the swimming speed. Figure 2 shows the change in the bacterial swimming speed. Berg et al. also determined the time of tumble using this method [7]. Although our data is a two-dimensional projection of three-dimensional motion, a decrease in the swimming speed is seen at the time of tumble motion.

**Fig. 2.**
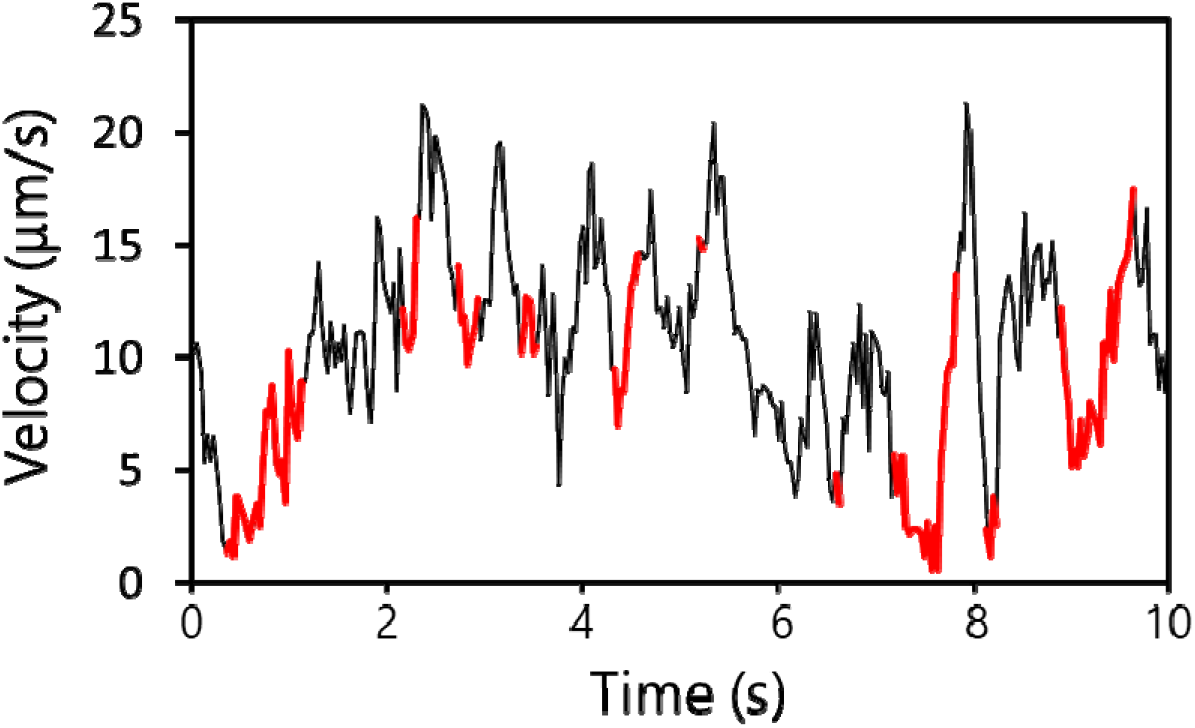
Relationship between the time of tumble and the decrease in the swimming velocity. The black and red lines correspond to the durations of runs and tumbles, respectively.

The characteristic time of the diffusion of L-serine can be roughly estimated from the diffusion constant *D*. In aqueous solution, the diffusion constant *D* is of the order 10^−9^ m^2^/s at 20°C. Consequently, the time *τ* required for L-serine molecules at the capillary tip to diffuse outside of the view can be approximately estimated using the below equation:

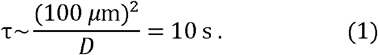

Hence this estimation implies that the concentration field of serine could be formed in several tens of seconds. To check the stable and concentric concentration field in the capillary assay mentioned, a fluorescent dye was used instead of L-serine. A capillary filled with 10 mM fluorescein (MW = 332, Wako, Japan) was observed using a laser scanning confocal microscope (FV10i, Olympus, Japan) with a 10× objective lens. The fluorescence distribution around the tip was almost concentric, which continued for at least 20 min. Thus, a similar concentration gradient of an attractant is also assumed in the observation of the chemotactic behavior of cells (within 15 min).

## III. Results

### 1. Trajectory of the trapped cell around an attractant

Among the 9 measured cells, data of 4 cells, the tracking time of which exceeded 100 s are especially shown in Fig. 3 (a). An example of a swimming trajectory (sample II in Fig. 3 (b)) is shown. The cell remains within 100 μm from the source of attractant. Whether this movement exhibits chemotaxis or the movement is only a random walk can be determined by calculating the mean square displacement (MSD). As shown in Fig. 3 (b), the MSDs of samples I-IV reach a plateau in the time region 10-100 s, indicating that the cells are attracted to the capillary tip. Without an attractant, the MSD should increase linearly with time.

**Fig. 3.**
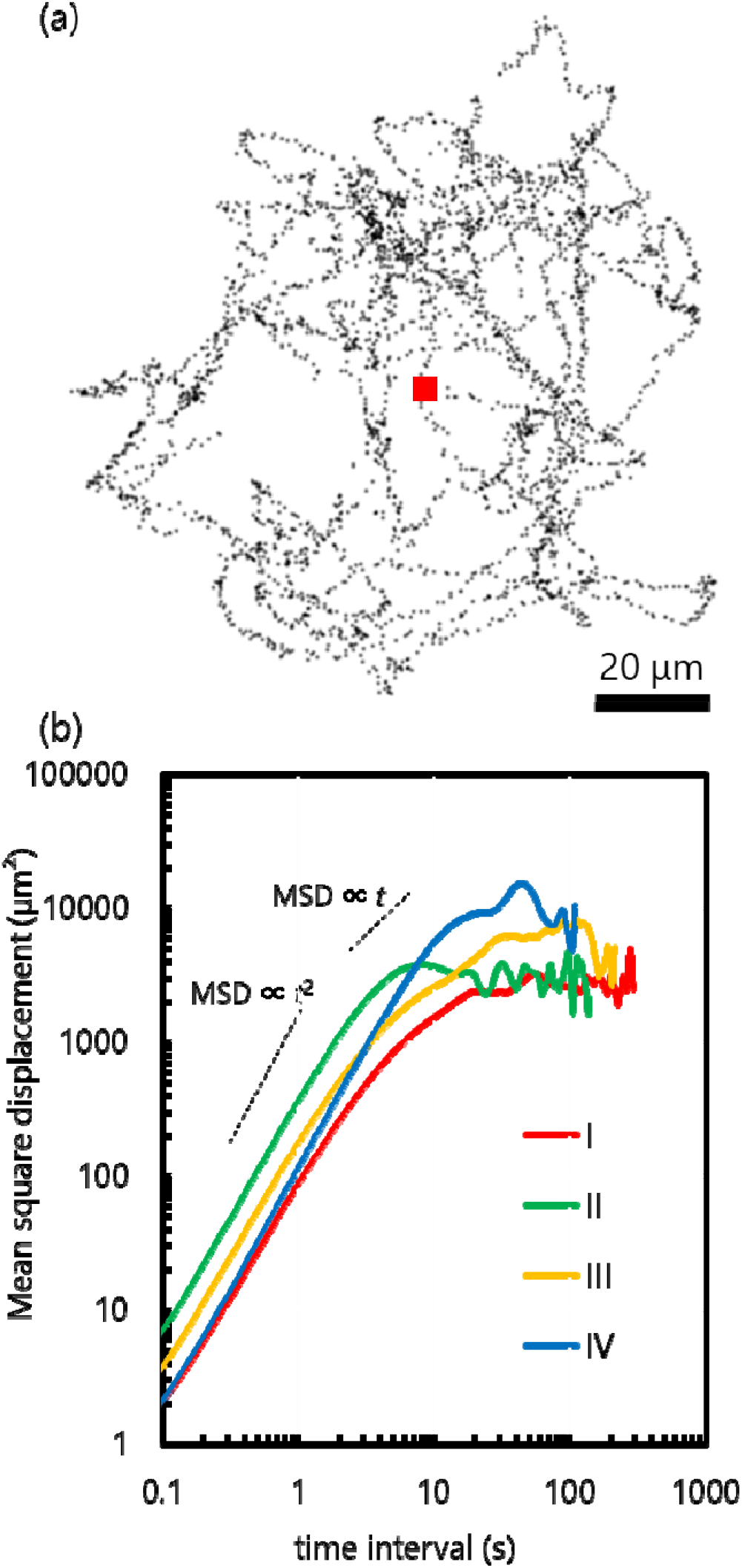
Single cell’s behavior around attractant source (L-serine). (a) Trajectory for sample II. The red square represents the position of the capillary tip. (b) Mean square displacement (MSD) of 4 cells.

### 2. Duration of runs

The duration of runs, the time between two adjacent tumbles, is one of the characteristics of bacterial chemotaxis, as reported in previous studies [7, 10]. In Fig. 4(a)(b), the duration of runs for 4 cells is shown, where runs are split, that is, with regard to swimming up and down the gradient. Data for the runs passing by the capillary tip (approximately 20% of total runs) are not shown. As shown in Fig. 4 (a), the average durations of runs “up” and “down” are roughly the same for all the 4 samples and there is no trend that runs are prolonged for cells that run “up”; this is in agreement with previous studies, where the duration of runs was measured by using many bacterial cells [7, 10], where duration for “up” movement is about 1.5 times longer than the duration of “down” movement. In Fig. 4 (b), the distribution of the duration of runs for “up” and “down” is shown. The vertical axis denotes the percentage of total runs for each cell. The short run (0-1 s) seems less frequent in “up” movement, but the shape of the distribution and peak position are almost the same. Long runs (2-3 s) are seen in both “up” and “down” movements, which may obscure the difference in the average values (around 1 s).

**Fig. 4.**
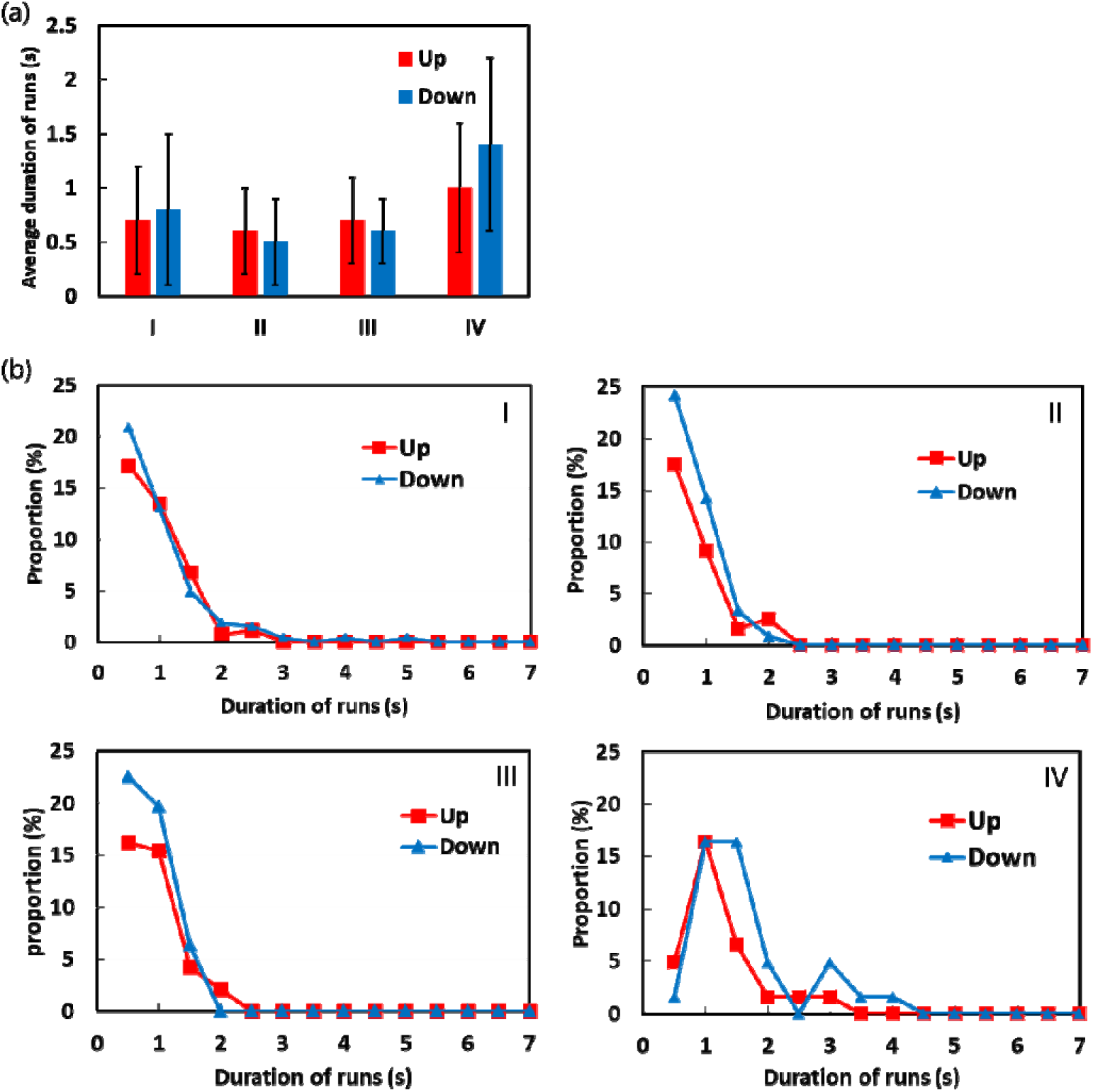
Duration of runs of a single cell around an attractant, (a) Comparison between swimming up and down the concentration gradient of the chemo-attractant, (b) Distribution of runs in samples I-IV. The vertical axis denotes the percentage of total runs for each cell.

### 3. Turn angle

The dependence of the swimming direction on the turning angle has been elucidated. As shown in Fig. 5 (a), the turn angle *θ* and the swimming direction *φ* are defined as the change in the swimming direction between two adjacent runs and the orientation angle relative to the capillary tip, respectively. In Fig. 5 (b), a plot of *θ* and *φ* for all 268 tumbles in sample I is shown. The orientation angle *φ* is less than 90° for “up” moving cells and larger than 90° for “down” moving cells. The turn angle *θ* is distributed uniformly when *φ* > 90°, whereas the distribution of *θ* is biased to a smaller angle when *φ* < 90°. This indicates that cells approaching the attractant source have a smaller average turn angle than receding cells. The mean values of the turn angles are different between “up” and “down” movements in the *t*-test with a significance level of 5%.

**Fig. 5.**
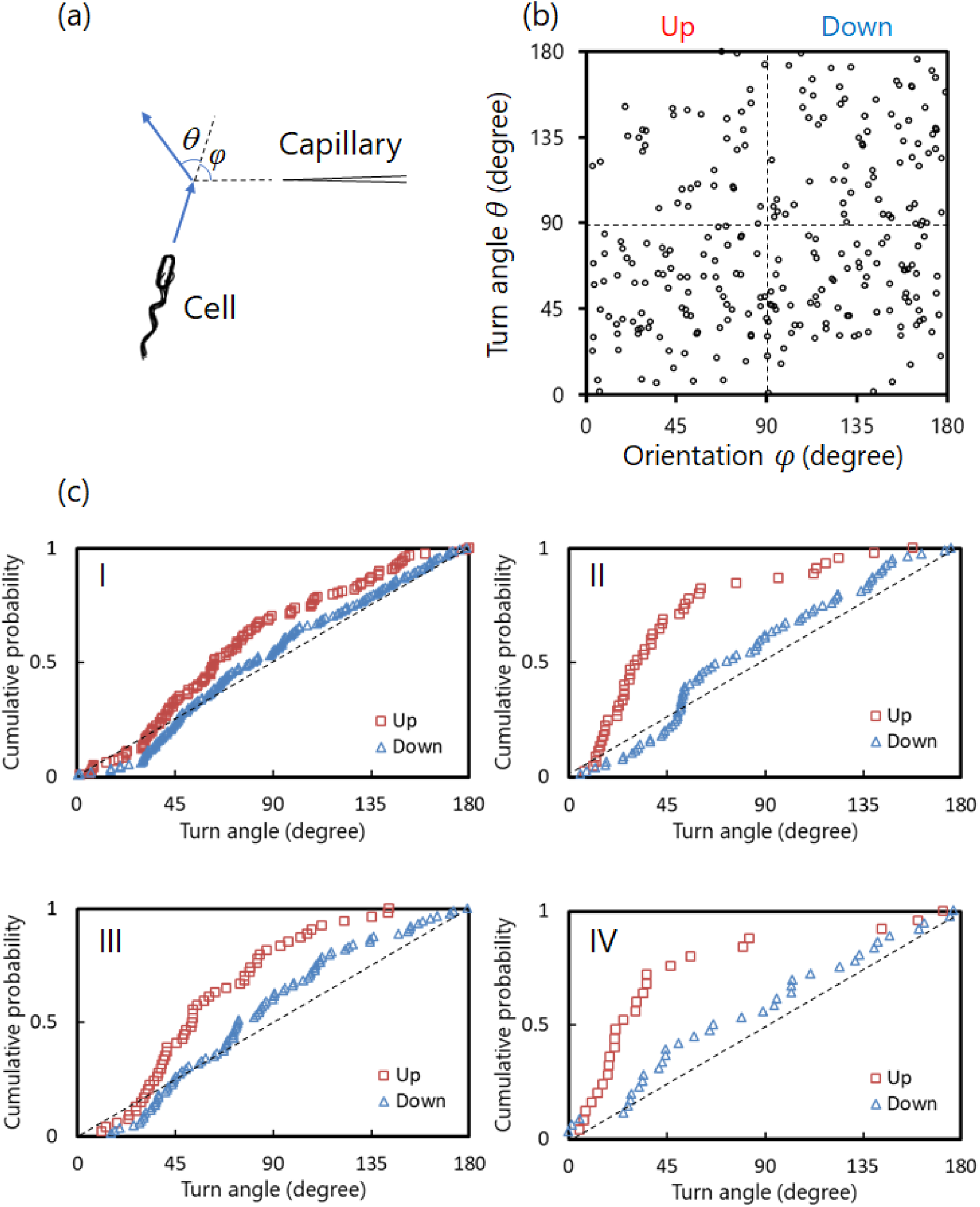
Turn angle distribution in a single cell, (a) definition of the directional change *θ* and the orientation *φ*, (b) *θ* – *φ* plot for sample I, and (c) cumulative probability of the turn angle.

As shown in Fig. 5(c), the bias can be clearly visualized by plotting the cumulative probability of *θ* for “up” (*φ* < 90°) and “down” (*φ* > 90°). The dashed straight line denotes that the turn angle *θ* has random distribution; *θ* is observed with equal probability at any angle. In samples I - IV, “down” is almost random and “up” has a plot above “down”, indicating that the directional change during the tumble after swimming up the gradient tends to be smaller. This tendency in the angle distribution was the same in the three-dimensional measurement using a piezoelectric-driven objective lens (see the supporting materials).

### 4. Biased random walk simulation

The effect of the observed bias in the turn angle on the degree of accumulation was investigated by numerical simulation. Goto et al. developed a mathematical model for bacterial chemotaxis based on the biased random walk [10, 15], where the cell changes the tumble frequency depending on the moving direction of the adjacent steps. In this subsection, we consider the bias in the turn angle in addition to the change in the tumble frequency.

We have herein provided the details of the mathematical model. As schematized in Fig. 6 (a), an attractant source is at the origin, and a concentric concentration field due to diffusion is assumed. The rules for the modeled cells are as follows:

- Cells migrate a constant distance *Δr* during a unit time step *Δt*.
- When the cell has moved toward the direction where the concentration of an attractant increases, that is, toward the source of an attractant, the cell moves in the same direction as the previous time step with the probability *α* (see Fig. 6 (a)). Alternatively, the cell changes its direction with probability 1 – *α*, where the direction in the next step follows the observed distribution “up” (Fig. 6(b)).
- A cell changes its direction in the random direction when the cell has moved toward the direction where the concentration decreases. The direction in the next step is determined randomly, reflecting the distribution “down” as in Fig. 5(c).

**Fig. 6.**
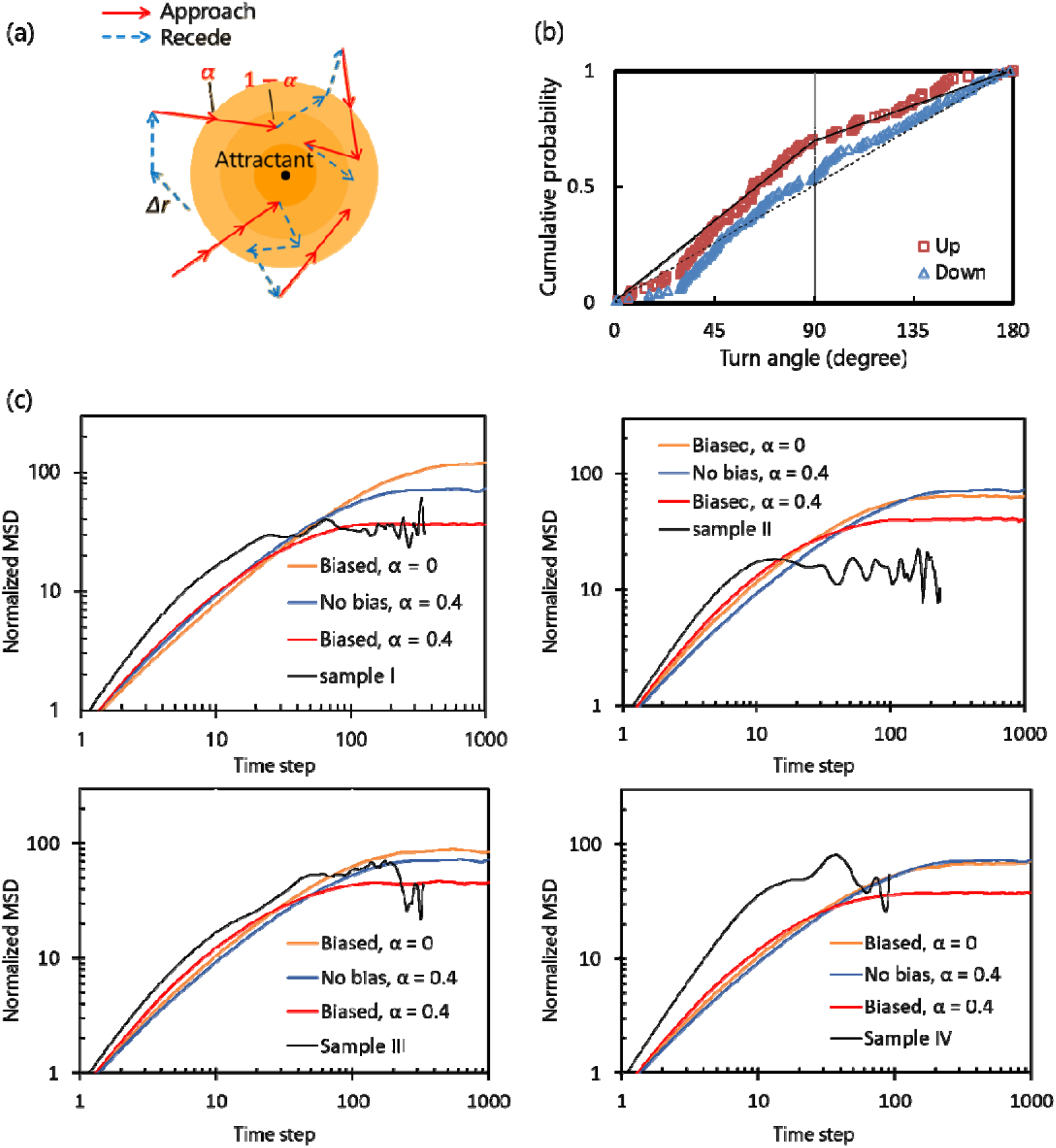
Biased random walk simulation of a swimming cell in the concentration gradient of a chemoattractant. (a) Scheme of the simulation model. Each cell migrates a constant distance *Δr* in a unit time step, corresponding to each arrow. Cells receding from the attractant source always tumble, whereas the behavior of cells approaching is stochastically determined with the probability *α*. (b) Modeling the difference in the turn angle between “up” and “down” for sample I. (c) Calculated mean square displacement (MSD) together with the observed one.

It must be noted that the swimming of cells up or down the gradient is dependent on the cell, and the cells do not sense how large the concentration gradient is. The parameter *α* (0 ≤ *α* ≤ 1) denotes the intensity of the chemotaxis; *α* = 1 is the maximum intensity and *α* = 0 is the minimum, that is, the random walk without biases. As for the turn angles, a simplified cumulative distribution of the observation was applied, as shown in Fig. 6(b). In the figure, the thick solid line represents a distribution for a cell that moved “up”, where the folding point at 90° implies that the probability that turn angle becomes smaller than 90° is higher. The turn angle for a cell that moved “down” is assumed to be a random value, as represented by a dashed line in the figure. Although the cumulative distribution is based on the two-dimensional measurement, we have confirmed the validity of the distribution by conducting three-dimensional measurement of turn angles. For details, see the supporting materials.

Fig. 6 (c) shows the calculated MSDs together with the observed ones. For comparison with the 2-dimensional microscopic observation, MSD was calculated from the 2-D projected trajectory. The MSD and time interval in the observation were made dimensionless by the average values of run length and duration of runs, respectively. The condition corresponding to the observation is “Biased, *α* = 0”. To compare the chemotaxis intensity in the case where cells change the duration of runs and have no bias in the turn angle distribution, calculation results with “No bias, *α* = 0.4” is also shown, which was the estimated *α* in a previous study [10]. Because both MSDs reach a plateau of same height, the observed bias in the turn angle distribution contributes to the bacterial chemotaxis.

As for the height of the plateau, the observed MSD (Samples I–IV) is several times smaller than “Biased, *α* = 0”. Considering *α* = 0.4, together with the biased angle distribution (“Biased, *α* = 0.4” in the graph), the plateau value of MSD becomes almost the same. However, the time interval until reaching the plateau is approximately 10 times smaller in the observation, and the slope of the MSD in the region where the time interval is of 1 – 10 steps is steeper in the observation. Thus, even in the calculation model considering the angle distribution and *α*, we were not able to reproduce the observed result; possible reasons will be described in the next section.

## IV. Discussions

### 1. Difference in MSD between observation and simulation

From Fig. 6 (c), it can be seen that the observed bias in the turning angle affects the bacterial behavior around an attractant. However, some parts of the calculated MSD curve are quantitatively not in correspondence with the observation results, the cause of which has been discussed in this subsection.

As can be seen from Fig. 4 (b), long runs of over 2 s is observed in both “up” and “down” movements; this could result in the slope of the observed MSD being steeper in the short-time region than in the simulation (Fig. 6 (c)). For the reason that the average duration of runs (~ 1 s) is adopted as the time step in the simulation, these long runs correspond to several time steps. Therefore, the slope of the MSD in the observation is almost 2 (ballistic) in the region of several time steps, whereas the slope is almost 1 (diffusive) in the simulation.

These long runs also affect the average value of the duration of runs: no difference between “up” and “down” that leads to the higher plateau of MSD in the simulation (“biased, *α* = 0” in Fig. 6 (c)). In Fig. 4 (b), runs of 1 s have a larger ratio for “Down”. Therefore it is considered that *α* would have a positive value more than 0. However, with regard to the average duration of runs including these long runs, the upward runs may not be longer than downward runs (Fig. 4 (a)).

One of the reasons for these long runs could be recovery from the adaptive state to the chemoattractant. Adaptation means that the bacteria become accustomed to the environment with the attractant and cannot detect the attractant unless bacterial cells move to a higher concentration. If the environment changes from no attractant to a uniform concentration (no concentration gradient) of the attractant, tumble frequency is reduced for the first minute, but then changes to the same frequency as the environment without the attractant [9]. In our experiment, it is possible that the bacteria migrated to a low-concentration region of the attractant (far from the capillary tip) and recovered from the adaptive state, which caused long runs independent of the swimming direction.

### 2. Duration of tumbles: guessing the number of flagellar filaments loosen

Although flagella were not observed in our experiment, the number of flagella (corresponding to the turn angle) loosened during the tumble can be inferred from the duration of the tumble. Switching of the rotational direction of each flagellar motor occurs independently, hence in a tumble with multiple flagella loosened, switching of multiple flagellar motors occurs within a short time, and the observed duration of the tumble is expected to be long. In Fig. 7, the distribution of the duration of tumbles for each swimming direction for 4 each cell is shown. The vertical axis is the ratio that becomes 1 when integrating only “up” or “down” movements (note the difference from Fig. 4). A large number of short-time tumble motions (~ 0.1 s) was observed in “up”, implying that the number of flagella loosened during tumble tends to be small in “up”.

**Fig. 7.**
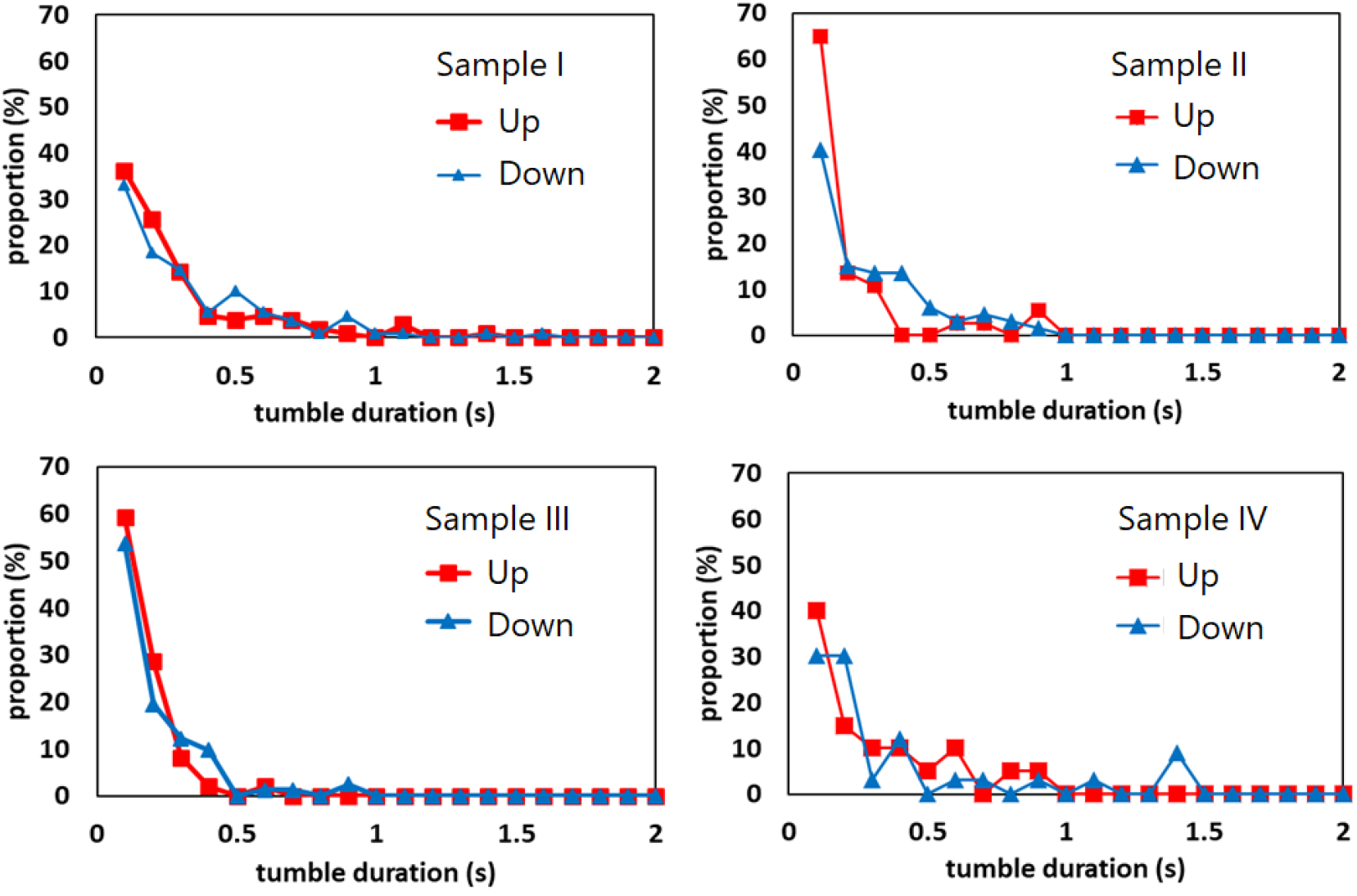
Distribution of the duration of the tumbles for samples I-IV.

## V. Conclusion

We investigated the turn angle in tumbles of peritrichous bacteria swimming across the concentration gradient of a chemoattractant. As predicted in Refs. 8, 13, and 14, we found that the turn angle depends on the concentration gradient that the cell senses just before the tumble. The turn angle is biased toward a smaller value when the cells swim up the concentration gradient, whereas the distribution of the angle is almost uniform (random direction) when the cells swim down the gradient; this behavior is appropriate because the nature of bacterial chemotaxis changes the switching rate of the rotational direction of the flagellar motors according to the environment. Cells swimming upward reduce the turn angle by loosening fewer flagellar filaments from the bundle during tumbles, resulting in accumulation at higher concentrations of the chemoattractant. The effect of the observed bias in the turn angle on the degree of chemotaxis was investigated by random walk simulation. In the concentration field where attractants diffuse concentrically from the point source, we found that this angular distribution clearly affects the reduction of the mean square displacement of the cell, that is, the bias in the turn angle distribution contributes to chemotaxis in peritrichous bacteria.

## Author Contributions

T.N. and T.G. designed research. T.A. performed experiments and analyzed data. T.N. wrote the paper.

## Acknowledgements

This work was supported by JSPS Kakenhi Grant Numbers JP18K03950 and 19K04193.

## Supporting Materials

### Three dimensional measurement of the turn angles

For the reason that in Fig. 5(c), the depth of direction is not shown in the turn angle distribution (the obtained angles are different from the actual one), we performed three-dimensional measurements and verified the validity of the angle measurement by two-dimensional observation.

A piezo element (THK PRECISION, PFHW2020-400U-S) was installed between the objective lens (OLYMPUS, LUCPlanFLN40X) and revolver of the inverted microscope (OLYMPUS, IX73), and connected to the PC via the controller (THK PRECISION, NCS6111S). The objective lens was vertically oscillated (period *T* of 0.125 s and amplitude of 50 μm) by a sinusoidal voltage applied to the piezo while recording with a high-speed camera (Photron, FASTCAM SA-X2) at 2000 fps. Images where the cell was in focus were extracted from the obtained movie, and then the z-position of the cell was deduced. The moments of the tumbles were judged based on the swimming trajectory. The judgment based on the change in the swimming speed (see Fig.2) was not carried out because the time resolution of the three-dimensional measurement (approximately the order of *T*) is as coarse as the tumble duration (~ 0.1 s).

In Fig. S1, the cumulative distribution of turning angles for each swimming direction (approaching or moving away) is shown. The angle distribution “2D” calculated from the trajectory projected on the two-dimensional plane is also shown together with the angle distribution calculated from the three-dimensional trajectory “3D”. Although there is some difference in the distribution between 3D and 2D, it is considered that the angle distribution used for the random-walk simulation is not very different.

---

**Fig. S1.**
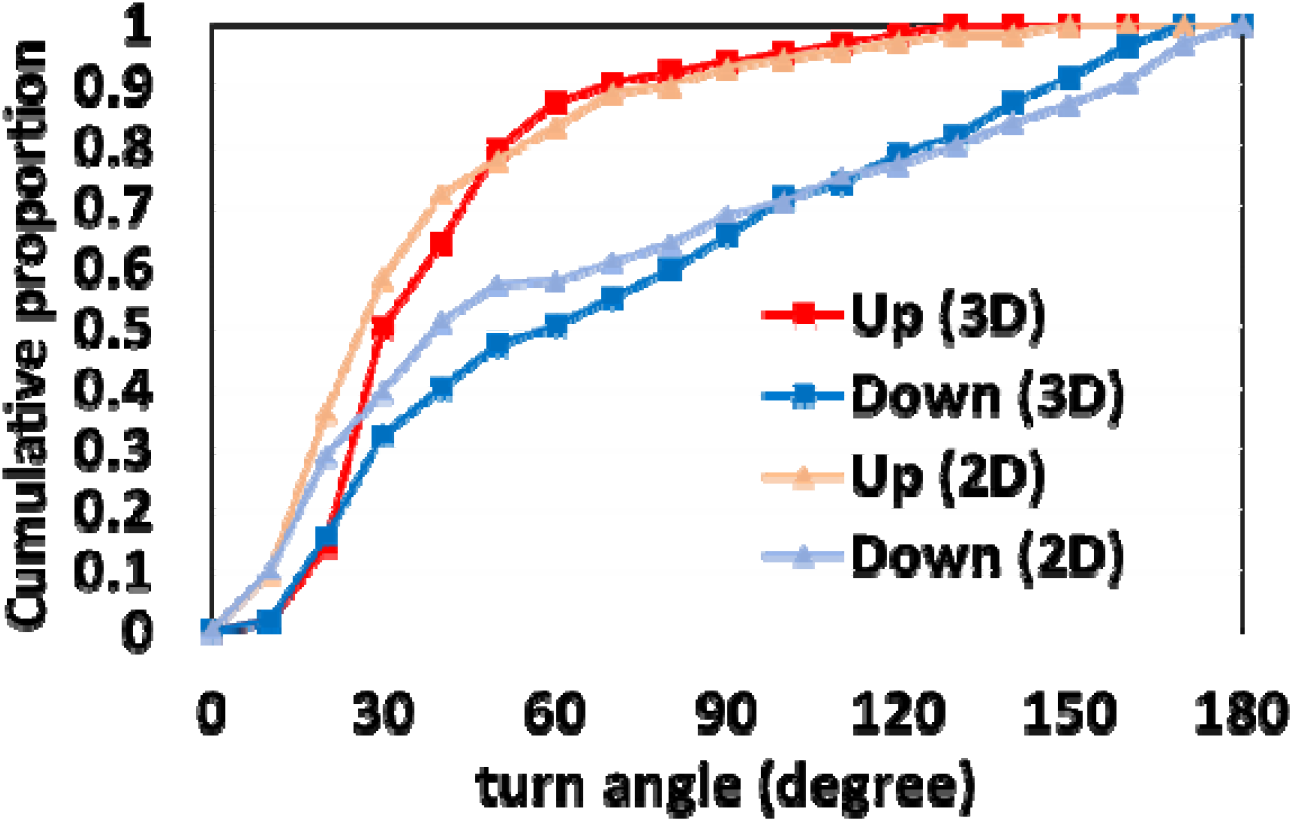
Cumulative distribution of the three-dimensionally measured turn angles. “2D” means the projection of the “3D”; the turn angles were deduced by using only the two dimensional coordinates (*x* and *y*) in the 3D measurement.

